# Functional interrogation of candidate *cis*-regulatory elements at the *LDLR* locus

**DOI:** 10.1101/2025.03.20.644385

**Authors:** Kyle Leix, Candilianne Serrano-Zayas, Sarah E. Graham, Brian T. Emmer

## Abstract

Regulation of *LDLR* gene expression plays an important role in the development of atherosclerotic diseases including heart attack and stroke. Although *LDLR* regulation by sterol response elements has been well characterized, the functional significance of other noncoding regions at the *LDLR* locus remains poorly defined. We developed and applied a high throughput CRISPR screen to test the functional significance of candidate *LDLR cis*-regulatory elements in their native genomic context. Analysis of our screen results revealed a discrete region in the first intron of *LDLR* with a significant impact on cellular LDL uptake. We validated the presence of enhancer activity in this region by confirming that its disruption reduced endogenous *LDLR* expression while its insertion upstream of a minimal promoter augmented reporter gene expression. We then applied a massively parallel reporter assay to fine map enhancer activity in this region to a 129 bp interval that is highly conserved among vertebrates, exhibits biochemical hallmarks of enhancer activity, is enriched for transcription factor binding motifs, and contains a common genetic variant (rs57217136) that has been associated with human LDL cholesterol levels by genome-wide association studies. Overall, these findings demonstrate the power of CRISPR screening to interrogate candidate CREs and support the functional significance of an enhancer in the first intron of *LDLR*.

## INTRODUCTION

The amount of LDL receptor expressed in the liver is a major determinant of a person’s lifetime risk of atherosclerotic cardiovascular diseases (ASCVD) including heart attack and stroke. Whereas genetic deficiency of *LDLR* causes familial hypercholesterolemia and early onset ASCVD, genetic variants associated with increased LDLR activity are protective against ASCVD^1,2^. Likewise, multiple classes of therapeutics that augment hepatic LDLR activity are effective in preventing the development of ASCVD^3^.

Pioneering studies of *LDLR* gene regulation have firmly established the importance of SREBP transcription factors and their binding to sterol response elements (SREs) in the promoter of the *LDLR* gene^4–6^. However, eukaryotic gene regulation is a complex process in which the activity of individual transcription factors may be modulated by interactions between promoters and *cis*-regulatory elements (CREs) of noncoding genomic DNA. The total number of CREs in the human genome is estimated to be on the order of hundreds of thousands to millions^7^. For *LDLR*, several lines of evidence point toward a complexity of regulation beyond SREBP binding alone. First, different genes containing SREs in their promoters vary widely in relative expression^4^. Second, other transcription factors that modify *LDLR* expression through sterol-independent mechanisms have been identified^4,8^, including in CRISPR screens by us^9^ and others^10,11^. Third, human genome-wide association studies have identified several noncoding common genetic variants associated with LDL cholesterol that reside near the *LDLR* locus but do not localize directly to SREs^12–14^; a targeted study of 4 of these detected a functional influence for 2^15^. Finally, genome-wide profiling experiments have identified candidate CREs at the *LDLR* locus that do not colocalize with SREs^16–19^. Collectively, these observations support the likelihood that *LDLR* expression is influenced by currently unrecognized CREs and their associated DNA-binding proteins.

We previously reported a high-throughput CRISPR screen of the coding genome for *trans*-regulators of cellular LDL uptake^9^. The rigor of our screening pipeline was strongly supported by our clear detection of known regulators of *LDLR* expression as well as our validation and mechanistic investigation of select novel regulators^9,20,21^. In this study, we now report our adaptation of this same CRISPR screening approach to interrogate the noncoding landscape of the *LDLR* locus, leading to our identification and subsequent fine mapping of an enhancer in the first intron of *LDLR*.

## RESULTS

### Design and synthesis of a custom CRIPSR library targeting putative CREs at the *LDLR* locus

To nominate candidate CREs for functional testing, we identified target regions based on multiple data sets. We first analyzed associations of common genetic variants with circulating LDL cholesterol levels in a recent multi-ancestry GWAS of over 1.6 million individuals^14^, finding a cluster of 73 strongly associated variants that localized within a ∼25 kb region around the *LDLR* transcription start site (TSS) and exhibited a high degree of linkage disequilibrium with each other; we included this entire interval in our target set (Supplemental Table 1, Fig S1A). Additionally, we performed a conditional analysis on the lead SNP in this region (rs73015024) to identify another 29 variants within 500 kb of the *LDLR* TSS that were independently associated with LDL cholesterol in individuals of European ancestry at genome-wide significance; we included 1 kb intervals centered on each of these 29 SNPs (Supplemental Table 2, Fig S1B). We also analyzed publicly available data sets^17–19,22^ to identify all regions within 200 kb of the *LDLR* TSS that exhibited biochemical features predictive of *cis*-regulatory activity, including an open chromatin state (ATAC-seq and DHS-seq peaks) and/or proximity to histone marks associated with enhancer activity (H3K27ac ChIP-seq peaks) (Fig S2). In this same interval we also identified regions with a high degree of evolutionary conservation among primates. Finally, we included all regions in this interval that were predicted by the ENCODE project to exhibit *cis*-regulatory activity^22^. In total, we identified 12,375 gRNA targets over an aggregate ∼70 kb of noncoding genomic space at the *LDLR* locus (Supplemental Table 3). As internal negative controls, we added 100 non-targeting gRNAs and 10 gRNAs targeting the AAVS1 ‘safe harbor’^23^. As internal positive controls, we added 10 experimentally validated gRNAs apiece targeting the coding sequences of *LDLR* or its negative regulator *MYLIP*^9^. We then synthesized a CRISPR library containing these gRNA sequences, verified library representation by deep sequencing, and generated and titered lentiviral stocks for pooled mutagenesis of target cell lines.

### Interrogation of putative *LDLR* CREs by CRISPR screening

To functionally interrogate putative *LDLR* CREs, we next tested our CRISPR library in HuH7 cells (Fig 1A). This cell line has been established as a model of hepatocyte LDL catabolism^24^ and, in our prior screen of the coding genome, exhibited clear dependence of LDL uptake on canonical regulators of *LDLR* expression including *SCAP*, *MBTPS1*, *MBTPS2*, and *MYLIP*^9^. For each of 4 independent biologic replicates, we transduced ∼25 million HuH7 cells at a multiplicity of infection of approximately 0.5, such that most individual cells would not receive multiple gRNAs. After 2 weeks of passaging to allow for target site editing, we incubated pools of transduced cells with exogenous fluorescently conjugated LDL, detached cells, and performed flow cytometry to collect 10% subpopulations of edited cells with the greatest and least amount of LDL uptake. We then quantified the abundance of each gRNA by massively parallel sequencing and assessed for enrichment between selected populations (Supplemental Table 4).

**Figure 1.**
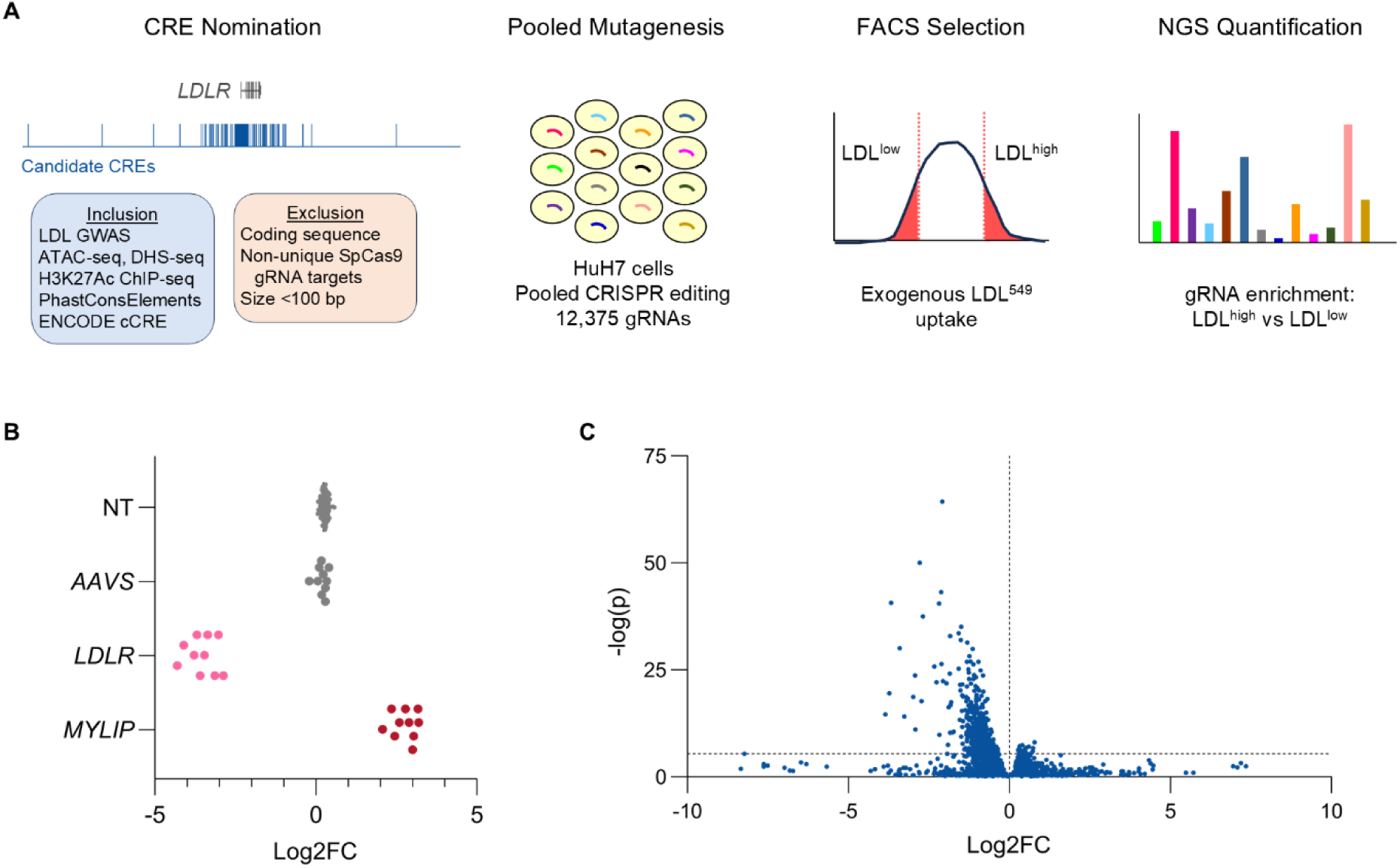
CRISPR screen for noncoding regulators of *LDLR* expression. (A) Schematic overview of CRISPR screen design with bioinformatic selection of candidate CREs and their pooled targeting by CRISPR indel formation followed by FACS isolation of individual cells with aberrant LDLR activity and quantification of enriched gRNAs by next-generation sequencing (NGS). (B) Segregation between LDL^high^ and LDL^low^ populations of internal control gRNAs either lacking a genomic target sequence (NT) or targeting the *AAVS* safe harbor (both in gray), or targeting the coding sequences of *LDLR* (pink) or its negative regulator *MYLIP* (dark red). (C) Volcano plot of individual gRNAs targeting *LDLR* candidate *cis*-regulatory elements, with x-axis representing log2 fold-change in LDL^high^ cells relative to LDL^low^ cells and y-axis representing the negative log p-value as determined by DESeq2. Dashed lines indicate log2 fold-change of 0 and the p-value significance threshold (4.08×10^-6^) as determined by Bonferroni correction of *p* = 0.05 for 12,245 tests. Control gRNAs displayed in (B) are omitted from (C).

We examined the technical performance of our screening pipeline by assessing the segregation of control gRNAs. As expected, gRNAs targeting the coding sequence of *LDLR* and *MYLIP* were consistently depleted or enriched, respectively, in LDL^high^ cells relative to LDL^low^ cells, whereas gRNAs lacking a genomic target site or targeting the *AAVS1* locus were similarly represented in either population (Fig 1B). The segregation of these control gRNAs was reproducible in pairwise comparisons of independent biologic replicates (Pearson coefficients 0.86-0.96, Fig S3).

Of the remaining gRNAs targeting putative *LDLR* CREs, the majority showed no significant difference between LDL^low^ and LDL^high^ populations (Fig 1C, Supplemental Table 4). Aside from 10/10 control *MYLIP*-targeting gRNAs, we observed few individual gRNAs with significant enrichment in LDL^high^ populations, suggestive of a lack of repressive elements among the regions targeted by our library. In contrast, we found 627 gRNAs to exhibit significant depletion (Bonferroni-corrected p-value < 0.05) in LDL^high^ cells, consistent with indel formation at their target sites leading to a decrease in *LDLR* expression. As expected, the strongest effects were observed for gRNAs predicted to generate double-strand breaks near intron-exon junctions (Fig S4), likely due to a disruption of the coding sequence and/or mRNA splicing by the resulting indels.

### Spatial organization of functionally significant *LDLR* gRNA targets

To identify genomic regions important for *LDLR* expression, we next plotted the enrichment scores for each gRNA in our screen by the genomic coordinate of its target site. Visualization of the 30 kb interval containing the cluster of LDL GWAS-associated common genetic variants revealed a striking spatial distribution of functional activity in our screen (Fig 2); whereas the entire region upstream of the *LDLR* promoter showed minimal activity in our screen, many gRNAs targeting the first *LDLR* intron were significantly enriched in LDL^low^ cells, suggesting that indel formation at their target sites caused a reduction in *LDLR* expression. While some of the active gRNAs targeted sequences near the first intron-exon junction, most targeted regions more distal from the first exon including a cluster of activity in a ∼0.6 kb region >500 bp away from the first intron-exon junction. We did not directly interrogate the *LDLR* promoter, as its overlap with an annotated antisense RNA led to its filtering from our candidate list.

**Figure 2.**
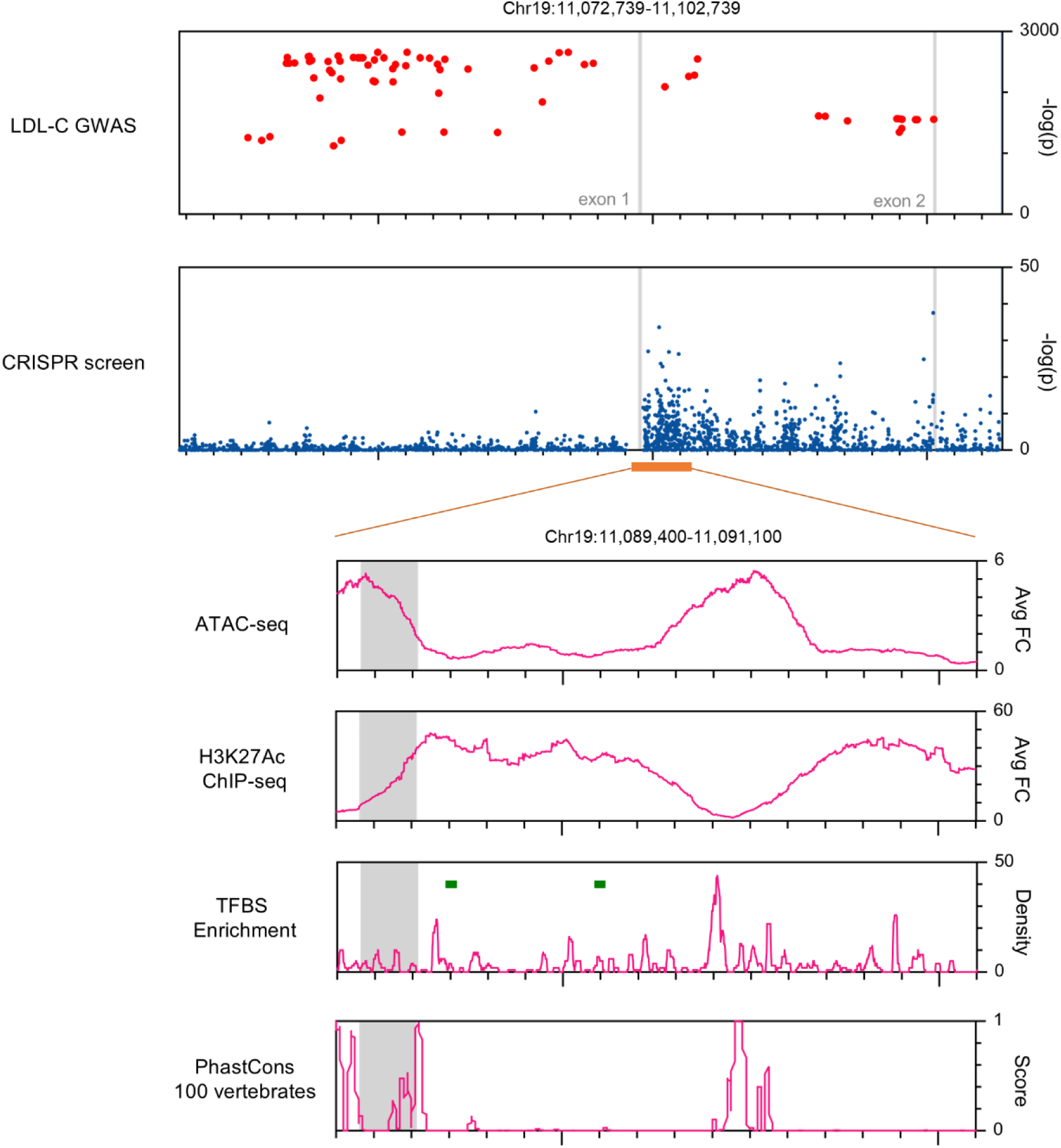
Spatial organization of CRISPR screen results. Candidate 30 kb interval spanning all common variants associated with LDL cholesterol with *p* < 10^-1000^ by GWAS (visualized in red) aligned with CRISPR screen results for the negative log p-value for depletion in LDL^high^ relative to LDL^low^ cells of each individual gRNA in the screen arranged by the coordinate of its predicted double-strand break site. Associated tracks for the 1.7 kb interval with maximum activity in the screen are displayed for human liver tissue ATAC-seq and H3K27Ac ChIP-seq, as well as the density of transcription factor binding motifs identified by JASPAR in each region and evolutionary conservation scores across 100 vertebrates. Exons are shaded in gray. Green boxes on the TFBS plot correspond to canonical SREBP binding motifs. CRISPR screen source data is provided in Supplemental Table 4.

We cross-referenced this functional map of the *LDLR* early first intron with other data sets that might further suggest *cis*-regulatory activity (Fig 2). Indeed, we found this cluster of targets for functionally significant gRNAs to colocalize with open chromatin as indicated by ATAC-seq of human liver tissue. This region also contained a distinctive enhancer-associated “peak-valley-peak” distribution for histone H3K27-acetylation^25^ in human liver tissue as well as an enrichment in JASPAR-predicted transcription factor binding motifs^26^. Included among the transcription factor binding motifs were two sites for SREBP binding, though most gRNAs that affected LDL uptake in this region targeted sites >100 bp from this motif. This region was also highly conserved among vertebrates, with PhastConsP scores approximating those observed for the nearby coding sequence of exon 1. Collectively, these findings suggest the presence of a conserved enhancer in the first *LDLR* intron that operates independently of direct SREBP binding.

### Validation of enhancer activity in the first intron of *LDLR*

To confirm the functional significance of the first intronic *LDLR* region to endogenous *LDLR* expression, we first selected 2 individual gRNAs targeting this region with highly significant depletion in LDL^high^ cells in our screen. We cloned these sequences into individual lentiCRISPRv2 constructs, transduced and selected HuH7 cells, and performed phenotypic characterization of the resulting polyclonal populations. In comparison to control cells transduced with a non-targeting gRNA, cells with targeting of the *LDLR* first intronic region exhibited significant reductions in LDL uptake (Fig 3A) and *LDLR* mRNA levels (Fig 3B). Transduction of these constructs into HepG2 cells likewise resulted in a reduction in baseline LDL uptake, though this was more apparent after culturing cells in lipoprotein-depleted conditions (Fig 3C).

**Figure 3.**
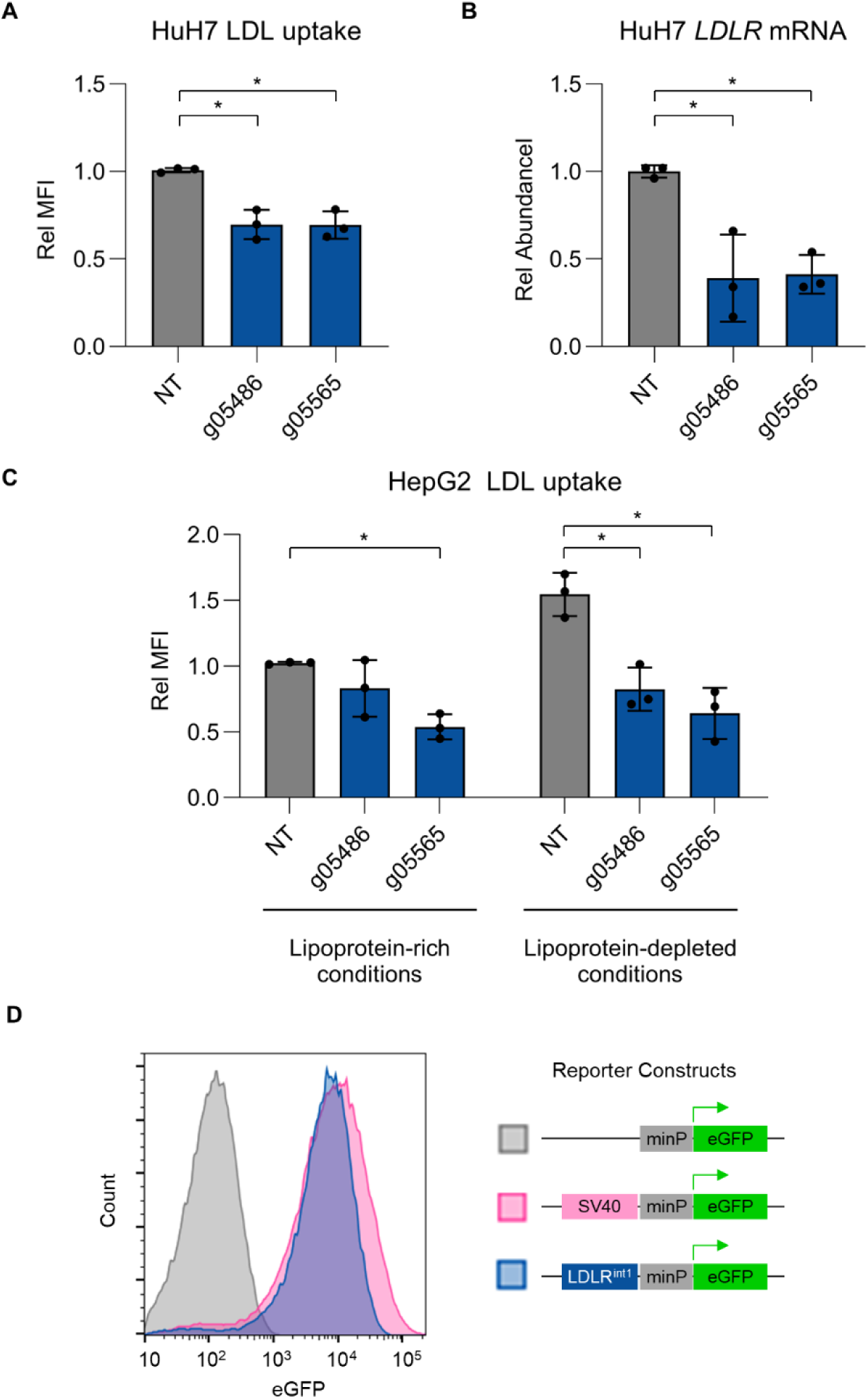
Validation of enhancer activity in the first *LDLR* intron. (A) Relative mean fluorescence intensity for LDL uptake by HuH7 cells transduced with a control nontargeting gRNA (gray) or individual gRNAs targeting the first *LDLR* intron that showed highly significant enrichment in the CRISPR screen (blue). (B) Relative *LDLR* mRNA levels as determined by qRT-PCR of mRNA preparations from the same HuH7 cells depicted in (A). (C) Relative mean fluorescence intensity for LDL uptake by HepG2 cells transduced with the same individual gRNAs as in (A-B) and cultured in the presence of normal serum or lipoprotein-depleted serum for 48 hr prior to the LDL uptake assay. (D) Schematic representation of lentiviral reporter constructs containing an eGFP coding sequence with a minimal promoter and an upstream test sequence. Histograms depict eGFP fluorescence for cells transduced with each construct in parallel at equal multiplicity of infection. For all panels, individual data points represent independent biologic replicates and asterisks indicate *p* < 0.05 for the indicated sample relative to control cells transduced with a nontargeting gRNA by one-way ANOVA with Dunnett’s correction for multiple hypothesis testing.

To further test for conventional enhancer activity in this region, we next PCR-amplified 567 bp of HuH7 genomic DNA corresponding to the region with maximum activity in our screen and inserted this sequence upstream of a minimal promoter and eGFP coding sequence in a lentiviral reporter plasmid^27^. Analysis by flow cytometry revealed the *LDLR* first intronic sequence to confer a ∼50-fold upregulation of eGFP expression relative to the parental construct (Fig 3D). The magnitude of effect for the *LDLR* first intronic sequence was comparable to that observed for the positive control, a SV40 sequence that is well-characterized to exhibit strong enhancer activity^28,29^. Together with the individual gRNA validation described above, these findings confirm the presence of enhancer activity within the first intronic region identified by our CRISPR screen.

### Fine mapping of enhancer activity in the first intron of *LDLR*

Although our CRISPR screen revealed enhancer activity within the first intron of *LDLR*, interpretation of its precise localization was complicated by intrinsic limitations of the approach, including the irregularity of SpCas9 gRNA targets across the region, the expected variability in activity for different gRNAs within the pool, and the expected heterogeneity of indel sizes arising at target sites. Therefore, to fine map enhancer activity in this region we instead developed a massively parallel reporter assay (MPRA). In this approach, large pools of candidate *cis*-regulatory sequences are tested in parallel for their ability to augment expression of a reporter gene^30^. We designed an oligonucleotide pool with 74 overlapping tiles of variable sizes (172, 129, 86, and 43 bp) that spanned an 860 bp interval with the most significant activity in our CRISPR screen (Fig 4A, Supplemental Table 5). We designed our library to represent the major allele in gnomAD^31^, notably including a common single nucleotide insertion (rs59281581) that has an estimated allele frequency of 99.8% but is not present in the hg38 reference sequence. We also included 3 positive and 3 negative internal control sequences that did or did not demonstrate *cis*-regulatory activity in a prior MPRA in HepG2 cells^32^. We amplified these sequences from oligonucleotide pools and cloned them into the pLS-SceI reporter backbone with incorporation of a random 15N barcode in the 5’ UTR of each construct. Sequencing of the resulting plasmid pool confirmed the synthesis of 69/74 expected tiles with a median of 2653 independent 5’UTR barcodes for each (Fig S5A, Supplemental Table 6). Expected tiles that were absent all shared the same 86 bp sequence with 78% GC content and a predicted hairpin structure that likely precluded its efficient synthesis, amplification, and insertion into the reporter plasmid (Fig S5B).

**Figure 4.**
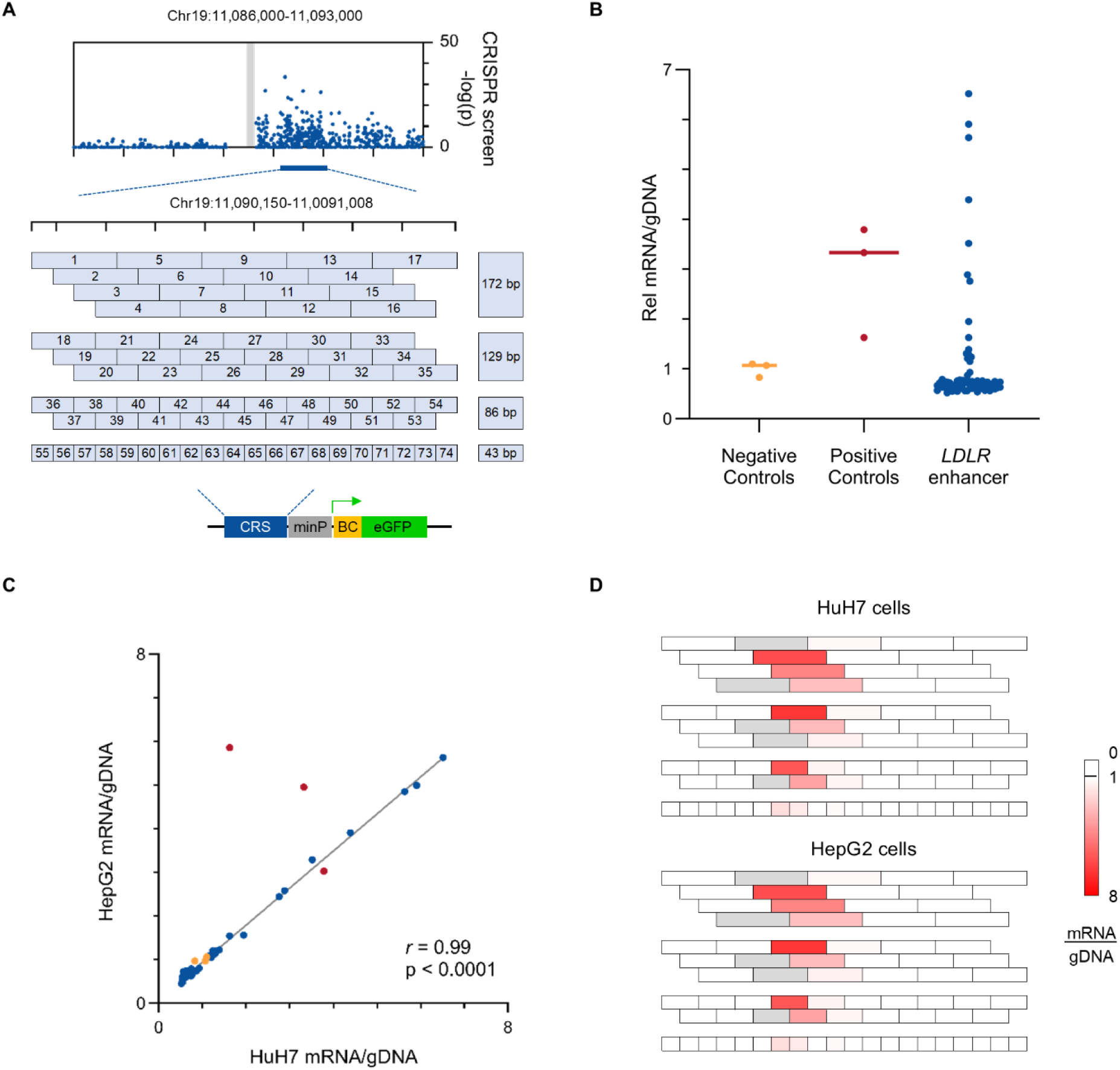
Fine mapping enhancer activity with a massively parallel reporter assay. (A) Design of the MPRA library containing overlapping tiles of noncoding sequence spanning different lengths of the candidate enhancer interval. (B) Relative reporter expression in HuH7 cells associated with each tile, derived from mapped sequencing reads of cDNA relative to genomic DNA. (C) Correlation of reporter expression conferred by each sequence in the MPRA library in HuH7 cells and HepG2 cells. Red and orange data points represent internal positive and negative controls, respectively, and blue data points represent candidate *LDLR* enhancer tiles. (D) Spatial organization of enhancer activity in each MPRA. A heat map is displayed for each tile visualized in (A), with greater enhancer activity represented by a deeper shade of red. Tiles with inadequate sequencing coverage in the plasmid pool are shaded in gray.

To functionally interrogate each tile, we then transduced HuH7 cells with the MPRA library, harvested genomic DNA and mRNA 2 days later, and performed massively parallel sequencing to quantify 5’UTR barcode abundance in each sample (Supplemental Table 7). Analysis of cDNA/gDNA ratios confirmed an increase in expression for positive controls relative to negative controls (Fig 4B, Supplemental Table 8). Of the other 69 test sequences, most exhibited no significant increase in activity relative to negative controls. A subset of test sequences however did show robust activity comparable to or exceeding that of the positive control sequences. To examine the generalizability of these findings across cell lines, we repeated the same MPRA approach in HepG2 cells, again finding discrimination of control sequences and a very high degree of concordance for the activity of test tiles between HuH7 and HepG2 cells (Fig 4C, *r* = 0.99, *p* <0.001).

Strikingly, the individual tiles that exhibited the greatest enhancer activity in our HuH7 and HepG2 MPRA studies all overlapped the same region (Fig 4D). This region was approximately centered on the ATAC-seq peak and the valley of the H3K27 acetylation ChIP-seq peak-valley-peak in this region, was highly conserved among vertebrates, and contained several motifs for transcription factor binding as well as one of the common genetic variants (rs57217136) strongly associated with LDL cholesterol by GWAS^12–14^ (Fig S6).

### Validation of fine-mapped enhancer activity

To validate the presence of enhancer activity in the narrower region identified by our MPRA studies, we next generated reporter constructs harboring 3 different fragments of the putative fine-mapped enhancer in both the forward and reverse orientations (Fig 5A). For 2 of the 3 fragments (containing tiles 6 and 42), we found enhancer activity to fully recapitulate that of the original full length construct; in each case, this activity was observed in either orientation, consistent with classical enhancer activity. For the 3^rd^ and largest fragment containing tiles 5 through 8, enhancer activity was present in either orientation though partially reduced in the reverse orientation, potentially related to an increased distance between the core active sequence and the minimal promoter.

**Figure 5.**
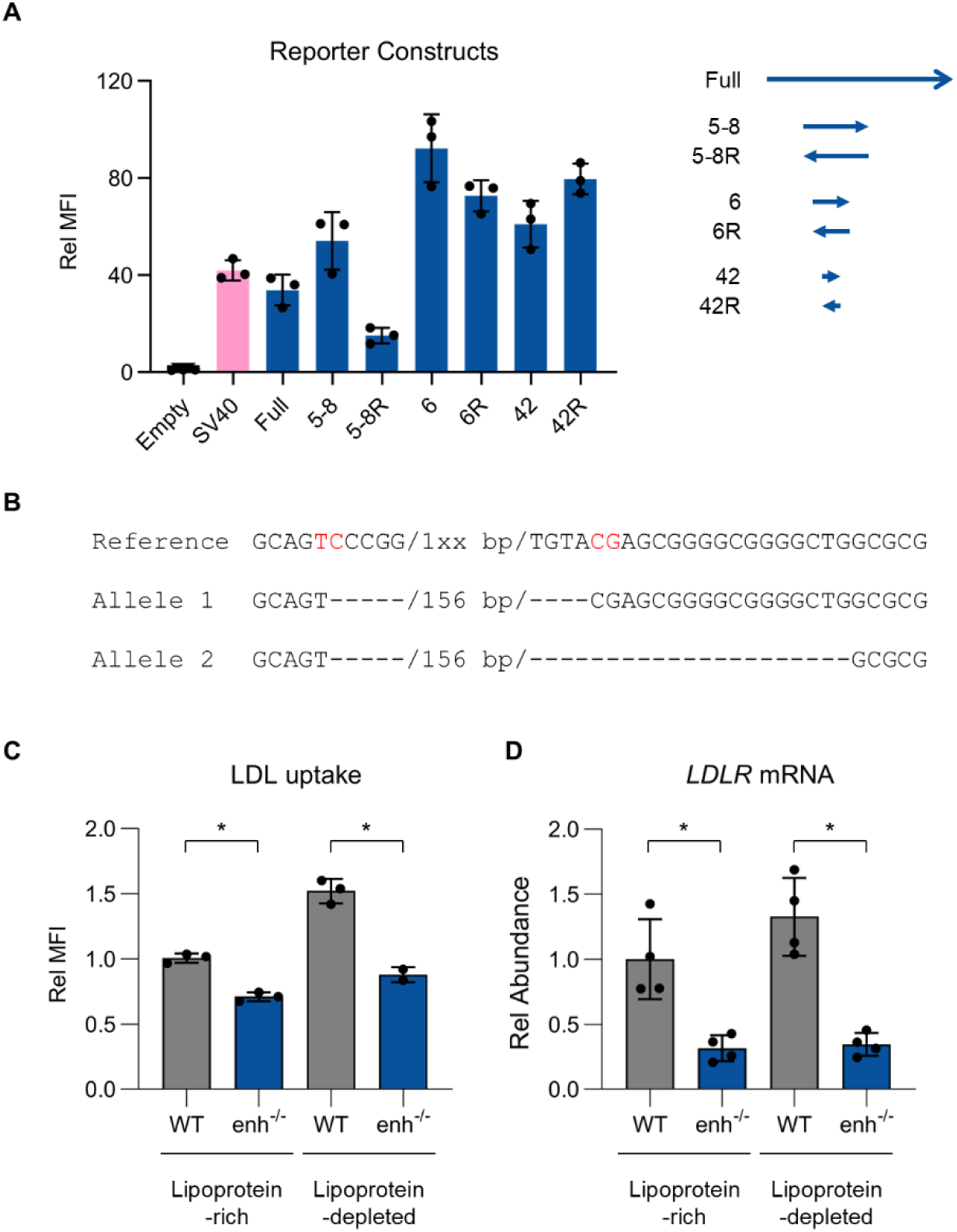
Validation of fine-mapped enhancer activity. (A) Mean fluorescence intensity (MFI) of eGFP relative to empty vector control for individual reporter constructs containing either the full length region containing the *LDLR* enhancer or the individual fragments identified by the MPRA in the forward or reverse orientations. (B) Genotype of a single cell clone recovered after transfection with a pair of gRNAs predicted to generate double-strand breaks between the nucleotides highlighted in red. (C-D) Uptake of fluorescent LDL (C) and *LDLR* mRNA levels (D) for the enhancer-deleted clonal population relative to parental wild-type cells, following culture of each cell line in lipoprotein-rich or lipoprotein-depleted media. Individual data points represent independent biologic replicates and asterisks indicate *p* < 0.05 by Student’s t-test.

To test the functional significance of this fine mapped enhancer activity in the context of the endogenous *LDLR* locus, we next transduced HuH7 cells with constructs expressing SpCas9 and a pair of gRNAs whose target sites flanked the minimal tile containing complete enhancer activity by MPRA testing. Genotyping of resulting single cell clones identified a cell line with compound heterozygous loss of the fine-mapped enhancer in both alleles (Fig 5B). Phenotypic characterization of this cell line confirmed a defect in LDL uptake (Fig 5C) and a reduction in *LDLR* mRNA levels (Fig 5D) under both lipoprotein-rich and lipoprotein-depleted conditions. These findings are consistent with the results of the reporter assay testing and confirm the presence of an enhancer in the first intron of *LDLR*.

## DISCUSSION

Historically, the criteria for identifying *cis*-regulatory elements have evolved along with the tools available for their interrogation^7^. Early definitions of enhancers referred to the ability of DNA sequences to increase the expression of reporter plasmids^28,33^. Over time, descriptions of enhancers began to incorporate biochemical features such as hypersensitivity to DNAse treatment (reflecting a lack of nucleosomes), an association with distinctive histone marks, and the presence of transcription factor binding sites. Although each of these features are useful, they remain imprecise tools for CRE identification. Numerous counterexamples exist both for *bona fide* enhancers that lack classical biochemical features, and for DNA regions that were suspected to be enhancers based on these features but exhibited no such activity upon functional testing^7,34–37^. Furthermore, biochemical predictions of enhancer activity often depend on coarse signals (e.g. broad peaks) that preclude precise mapping to specific boundaries.

Recently, advances in genome engineering have revolutionized the field of functional genomics. While high-throughput CRISPR screening has been mainly applied to the coding genome, studies that have extended this approach to the noncoding genome have consistently demonstrated its power to define the functional significance of CREs in their native genomic context^38–43^. Consistent with these studies, we now report our identification of an enhancer in the *LDLR* first intron by high-throughput CRISPR screening in HuH7 cells. The fidelity of our results is strongly supported by our clear discrimination of control gRNAs, the reproducibility in gRNA enrichment across independent biologic replicates, the spatial clustering of our screen hits, and by the empiric validation of our top hit in follow up studies.

Intriguingly, we found that most noncoding regions of the *LDLR* locus with predicted CRE activity exhibited no impact on *LDLR* expression when targeted by CRISPR-mediated indel formation. This included the majority of the ∼25 kb interval spanning the common genetic variants with the strongest LDL cholesterol associations by GWAS^12–14^. This observation suggests that many of these variants may not directly influence *LDLR* expression but rather correlate with LDL due to their linkage disequilibrium with other causal variants. However, there are several caveats to this interpretation that warrant consideration. First, enhancer activity may be highly context-dependent and not recapitulated by the conditions of our screen; that is, a noncoding region may show no effect in our study yet still play an important role in *LDLR* expression in other cell types and/or in response to different developmental or environmental cues. Second, pooled CRISPR screening is inherently limited in detecting functionally redundant CREs that contribute to gene expression in aggregate but do not cause a discernible phenotype when disrupted individually. Third, our ability to disrupt candidate CREs is limited by the presence of SpCas9 PAM motifs (with an average density of 1 in every 8 bp across the human genome) in regions of interest. Similarly, individual gRNAs may vary substantially in their relative activities. A given CRE therefore may be missed by our approach if it exists in a region with a paucity of gRNA targets or with target sequences for gRNAs with low activity. Finally, the indels that arise at CRISPR target sites are heterogeneous among individual cells transduced with the same gRNA. For example, if a gRNA target site is adjacent to a given *LDLR* CRE, then only the subset of cells with indels large enough to disrupt this CRE may influence *LDLR* expression and drive enrichment of the gRNA in the selected population. These considerations are especially pertinent for any enhancers with modest effect sizes on *LDLR* expression that may fall below the power of detection of our approach.

These limitations notwithstanding, our study expands the functional understanding of the noncoding landscape at the *LDLR* locus. Most strikingly, we observed clear activity within the first intron of *LDLR* that we validated and fine mapped to a 129 bp region that was highly conserved, associated with biochemical hallmarks of enhancer activity, and enriched in transcription factor binding motifs. This region also contains a single common genetic variant (rs57217136) that is strongly associated with human LDL-cholesterol and in linkage disequilibrium with other top LDL-associated variants at the *LDLR* locus. Intriguingly, a prior study found the minor allele of rs57217136 to increase reporter gene expression in HuH7 cells, but in contrast to our results did not detect enhancer activity for the major allele^15^. The basis for this difference is unclear but may be related to details of experimental design including the reporter constructs and readouts used in each study.

The fine-mapped enhancer identified in our study does not contain a canonical sterol response element, the binding site for SREBP transcription factors and the primary established mechanism for *LDLR* gene regulation. Instead, this region contains putative binding sites for several transcription factors including: Krüppel-like factors (KLFs), some of which have been previously implicated in atherogenesis and hepatic metabolism^44,45^; specificity proteins (SP), which are evolutionarily related to KLFs and one of which (SP1) has been shown to act synergistically with SREBP at the *LDLR* promoter^46,47^; and glioma-associated oncogenes (GLI), which are involved in hedgehog signaling^48^. Although none of these transcription factors were identified in our prior LDL uptake screen of the coding genome^9^, functional redundancy among members of the same family could mask the development of a phenotype for any single gene disruption. A previous study also reported binding by RAR and STAT1 to this region for the minor allele of rs57217136, though no binding or enhancer activity was detected for the reference allele^15^. Further work will be necessary to dissect the relative contributions of these and other DNA-binding proteins to enhancer activity in the first *LDLR* intron.

## Acknowledgments

This research was supported by the National Institutes of Health K08-HL148552 (BTE), R01-HL167733 (BTE), R01-HL171013 (BTE), and the A. Alfred Taubman Medical Research Institute (BTE).

## Author Contributions

CSZ and BTE conceived the project. CSZ, KL, SEG, and BTE performed the experiments. All authors contributed to data analysis and interpretation. KL, CSZ, and BTE wrote the manuscript with input from all authors. All authors reviewed the final version of the manuscript.

## MATERIALS AND METHODS

### Cell lines and reagents

HuH7 and HepG2 cells (ATCC, Manassas VA) were cultured in DMEM supplemented with 10% fetal bovine serum, 10 U/mL penicillin, and 10 µg/mL streptomycin (Thermo Fisher Scientific, Waltham MA) in a humidified 5% CO2 chamber at 37°C. Cell lines were periodically tested for mycoplasma contamination and verified by microsatellite genotyping. Oligonucleotide sequences (Integrated DNA Technologies, Coralville IA) are provided in Supplemental Table 9. CRISPR constructs were generated by ligation of phosphorylated and annealed oligonucleotides into BsmBI-digested pLentiCRISPRv2^49^ (Addgene, Watertown MA, #52961) or pX459^50^ (Addgene #62988) as previously described^51^. Reporter constructs were generated by PCR amplification of the indicated noncoding sequence from genomic DNA or a plasmid template with primers designed to provide flanking homology arms for HiFi assembly into AgeI/SbfI-digested (New England Biolabs, Ipswich MA) pLS-SceI^27^ (Addgene #137725) as previously described^27^. A plasmid with the same backbone and a SV40 enhancer sequence was also obtained as a positive control (pLS-SV40-mP-EGFP, Addgene #137724). Plasmids were verified by Sanger sequencing or whole plasmid sequencing and lentiviral stocks were prepared in coordination with the University of Michigan Vector Core.

### Candidate CRE nomination and analysis

Summary statistics were downloaded from the Global Lipid Genetics Consortium multi-ancestry GWAS of LDL cholesterol levels in 1,654,960 individuals^52^. Individual SNP associations were visualized together with their linkage disequilibrium with the lead SNP using LocusZoom^53^. The 30 kb interval spanning all SNPs with p<10^-1000^ was selected for saturation editing. To identify more distal variants with independent associations with LDL levels, we performed a conditional analysis on the lead SNP (rs73015024) for all variants within 500 kb of the *LDLR* TSS with an allele frequency >1% in individuals of European ancestry using GCTA-COJO^54^; from this analysis we targeted a 1 kb interval centered on each of 29 SNPs independently associated with LDL. Additional candidate *LDLR* CREs were identified by analysis of genomic tracks downloaded from the ENCODE Project^22^ and UCSC Table Browser^55^ and are graphically depicted in Supplemental Figure 2. Identifying information for individual tracks is provided in Supplemental Table 10 and the basis for selection of individual gRNAs is provided in Supplemental Table 3. In brief, regions of open chromatin in hepatocytes were identified by colocalization with peaks for ATAC-seq or DHS-seq from human liver tissue and/or HepG2 cells^17,18^. Regions associated with histone marks of enhancer activity were identified by colocalization with peaks for H3K27Ac ChIP-seq from human liver tissue and/or HepG2 cells^19^. Evolutionarily conserved regions were identified by downloading the phastConsElements30way table from the 30 Primates track of UCSC Genome Browser, based on a previously described phylogenetic reconstruction^56^. Regions were excluded if they colocalized with exons in the NCBI RefSeq track of UCSC Table browser or if they spanned <100 bp of genomic space. All SpCas9 target sites within 500 MB of the *LDLR* TSS were downloaded from the CRISPR Targets track of USCS Table Browser and filtered for those whose predicted double strand break location overlapped the candidate CRE blocks identified above. For subsequent analyses of CRISPR screen results, additional data sets were downloaded from the ENCODE Project and USCS Table Browser. Predicted transcription factor binding sites were obtained from UCSC Table Browser track JASPAR TFBS, filtered for quality scores >=400, and enumerated at each chromosomal coordinate in the candidate enhancer interval. Evolutionary conservation was analyzed by downloading the PhastCons tables from the tracks for 30 Primates and 100 Vertebrates in UCSC Table Browser; rolling averages across a 11 bp window centered on each chromosomal coordinate were calculated for each table and are plotted in Fig 2.

### CRISPR library synthesis

A custom CRISPR library was synthesized as previously described^9,57^ to encode 12,375 unique gRNAs targeting the candidate CRE blocks identified above as well as 100 nontargeting negative control sequences, 10 negative control sequences targeting the *AAVS1* genomic “safe harbor”, and 10 positive control sequences apiece targeting the coding sequences of *LDLR* and *MYLIP* based on prior functional data^9^. Briefly, a pooled library of oligonucleotides was obtained that contained each gRNA sequence flanked by invariant annealing sites for PCR amplification (CustomArray, Bothell WA). This library was amplified using primers that provided additional flanks for HiFi assembly into BsmBI-digested pLentiCRISPRv2. Assembly products were electroporated into Endura electrocompetent cells (Lucigen, Middleton WI) and plated on 245 mm bioassay plates of LB-agar. Serial dilutions of electroporated cells were also prepared and colonies enumerated to confirm >50X library coverage and a >20:1 ratio for assembly reactions relative to control reactions with water in place of the oligonucleotide pool for initial PCR amplifications. Plasmid DNA was harvested from the resulting colonies and library representation was verified by massively parallel sequencing with an Illumina MiSeq single end 1×150 read and mapping individual reads to expected gRNA sequences using PoolQ as previously described^58^.

### CRISPR screen for modifiers of LDL uptake

For each of 4 independent biologic replicates, approximately ∼50 million HuH7 cells were distributed across 8 separate 15 cm diameter plates and transduced with the CRISPR lentiviral library at an estimated MOI of 0.5. Transduced cells were selected with 3 μg/mL puromycin beginning on day 1 post-transduction and continuing until control non-transduced cells were no longer viable. Selected cells were passaged every 2-3 days as needed, maintaining >1000X library coverage of total cell numbers with each passage. On day 14, cells were incubated with 4 µg/mL DyLight^549^-conjugated LDL (Cayman Chemical, Ann Arbor MI) in serum-free DMEM for 1 hr, harvested, and sorted by flow cytometry into subpopulations with the top and bottom 10% of LDL uptake. Genomic DNA was purified from each subpopulation and the integrated gRNA sequences were PCR amplified with incorporation of sample barcodes into the primers. Analysis of gRNA abundance between samples was performed by sequencing amplicons with an Illumina NextSeq paired end 2×100 read, mapping and demultiplexing reads with PoolQ, and assessing gRNA enrichment with Mageck^59^.

### Analysis of *LDLR* expression and LDL uptake

For comparison between control cells and cells with the indicated genetic perturbations, LDL uptake was assayed by incubating monolayers of cells in 6 well plates with 4 µg/mL of DyLight^549^-conjugated LDL in serum-free DMEM for 1-2 hrs at 37°C, washing with PBS, harvesting cells, and analyzing mean fluorescence intensity by flow cytometry. To adjust for the confounding influence of cell density on LDL uptake per cell, a standard curve was prepared by plating control cells at a range of densities and estimating cell density by the number of flow cytometry events for a fixed volume of cell suspension. LDL uptake was then normalized relative to the expected fluorescent intensity at a given cell density. For measurements of *LDLR* expression, cellular RNA was harvested with an RNEasy Plus Mini kit (Qiagen), converted to cDNA using Superscript III Reverse Transcriptase (Thermo Fisher), and analyzed by qPCR with Power SYBR Green (Thermo Fisher), gene-specific primers (Supplemental Table 9), and QuantStudio 5 (Thermo Fisher).

### Enhancer reporter assays

For initial testing of enhancer activity in the first *LDLR* intron, PCR was performed with PrimeSTAR GXL polymerase (Takara Bio, Kusatsu Japan) and the primers listed in Supplemental Table 9 to amplify the target region from HuH7 genomic DNA. Additional rounds of PCR were then used to append vector homology arms and incorporate a random 15N barcode into the 5’ UTR of the reporter construct. HiFi DNA assembly (New England Biolabs) was performed to insert the amplicon into AgeI/SbfI-digested pLS-SceI. Subsequent constructs were generated by similarly amplifying the region of interest from the parental enhancer construct. Lentiviral titers were calculated by transducing HuH7 cells at 20% confluence in 6 well plates, harvesting genomic DNA 2 days later, and performing qPCR to quantify the integrated lentiviral DNA relative to plasmid and host genomic DNA and translate these values to an estimated multiplicity of infection as previously described^27^. Transductions were then repeated with adjustment of each lentiviral stock volume to achieve the same MOI. Cells were harvested at 2 days post-transduction and analyzed for eGFP mean fluorescence intensity by flow cytometry.

### MPRA library synthesis

Individual overlapping tiles of varying sizes in the candidate region of *LDLR* intron 1 were selected for fine mapping as indicated in Supplemental Table 5. The overall strategy for library synthesis was performed as previously described^27^, with adaptations as indicated below. Briefly, an oligonucleotide pool was obtained (Integrated DNA Technologies) that included each candidate tile flanked by invariant sequences that served as annealing sites for amplification. Two rounds of PCR were performed with PrimeSTAR GXL polymerase using an annealing temperature of 60°C and the primers listed in Supplemental Table 9 that added additional flanks for HiFi assembly and incorporation of a random 15N barcode into the 5’ UTR of the reporter construct. The number of cycles was optimized by analysis of PCR amplicons on a Bioanalyzer DS1000 chip (Agilent Technologies, Santa Clara CA). Amplicons were purified with a QIAquick PCR Purification kit (Qiagen) and a total of 50 ng were combined with 200 ng of AgeI/SbfI-digested pLS-SceI for HiFi DNA assembly (New England Biolabs) at 50°C for 1 hr. The assembly products were then column purified and electroporated into Stbl4 *E. coli* cells (Thermo Fisher) using a MicroPulser (Bio-Rad Laboratories, Hercules CA) with setting Ec1. Electroporated cells were spread onto 245 mm LB-agar plates. Total colony counts were estimated by also plating serial dilutions of the electroporated cell suspension in parallel. Following verification of >100X colony counts relative to library size with a >20:1 ratio of colonies on assembly plates relative to control reactions containing digested vector alone, resulting colonies were scraped and plasmid DNA extracted using a Plasmid Plus Midi kit (Qiagen). The representation of the candidate tiles in the resulting plasmid pool was then tested by massively parallel sequencing. Amplicon libraries were prepared by 2 rounds of PCR using the primers indicated in Supplemental Table 9 to add flanking sample barcodes and sequencing adaptors. Optimization of PCR cycle number and assessment of amplicon library quality were again performed with a Bioanalyzer DS1000 chip. Sequencing was performed on an Illumina MiSeq instrument using the custom primers indicated in Supplemental Table 9. CRS sequences were extracted from reads 1 and 2 and associated with 15 bp CRS barcodes in index read 1. Individual barcodes were filtered to remove any with a read count <10 or with <99% association to the same unique CRS. The resulting array of barcode-CRS associations was used as a reference map for subsequent MPRA testing and is provided in Supplemental Table 6. For expected CRSs not detected in the plasmid library, a shared sequence was analyzed for predicted secondary structure by Unafold^60^. Lentiviral stocks were prepared from plasmid pools and titered by transducing HuH7 cells with a range of virus volumes and quantifying the resulting MOI by qPCR from genomic DNA with primers for integrated lentiviral sequence, plasmid backbone, and host genomic DNA as described above.

### Massively parallel reporter assay

HuH7 and HepG2 cells were seeded in 6 well plates at an estimated 20% confluence, with 5 wells used for independent transductions with the MPRA library at an estimated MOI of 5 and the remaining well serving as an uninfected control. Two days later, cells were harvested and split in half for extraction of genomic DNA and RNA using DNeasy and RNeasy Plus Mini kits, respectively (Qiagen). For each sample, 10 µg of RNA was treated with 10 units of Turbo DNAse (Thermo Fisher) for 30 mins at 37°C followed by addition of DNAse Inactivation Reagent for 5 mins at room temperature and conversion of 1.6 µg RNA to cDNA using the gene-specific primer P7-pLSmP-ass16UMI-gfp and Superscript III First-Strand Synthesis Kit (Thermo Fisher) per manufacturer’s instructions. Genomic DNA and cDNA samples were then used as templates for PCR amplification of CRS barcode sequences with primers providing additional barcodes for sample identity and Illumina adapters. The abundance of each CRS barcode in each sample was assessed by custom sequencing using the primers listed in Supplemental Table 9 on an Illumina NextSeq instrument. CRS barcode sequences were extracted from individual reads, filtered to remove any with <10 counts across all samples, demultiplexed by the sample barcode in index read 2, mapped to their associated CRSs, normalized to counts per million, and aggregated across CRS barcodes mapping to the same CRS. The relative expression conferred by each CRS was inferred by calculating the average ratio of normalized counts associated with each CRS in cDNA versus genomic DNA for each individual replicate. Normalized read counts for each CRS are provided in Supplemental Table 7 and expression analysis is provided in Supplemental Table 8.

### Enhancer deletion

HuH7 cells were co-transfected with an equal mixture of pX459 constructs harboring individual gRNAs targeting sequences upstream and downstream of the fine-mapped *LDLR* enhancer (Supplemental Table 9) using Lipofectamine LTX (Thermo Fisher) per manufacturer’s instructions. Puromycin selection was applied from days 1-3 post-transfection at which time no viable cells remained for control untransfected cells. Puromycin-resistant transfected cells were detached into suspension and diluted for single cell cloning into 96 well plates. Wells containing a single focus of cell proliferation were subsequently expanded and genotyped by extracting genomic DNA using a DNEasy kit (Qiagen), PCR amplifying the target region with PrimeSTAR GXL polymerase (Takara Bio) and primers flanking the predicted double-strand break site (Supplemental Table 9), performing amplicon sequencing (Plasmidsaurus, Eugene OR), and aligning the resulting reads using the Integrated Genomics Viewer^61^. A clone harboring compound heterozygous deletion of the enhancer was selected for subsequent analysis with LDL uptake and *LDLR* expression as described above.

---

**Figure S1.**
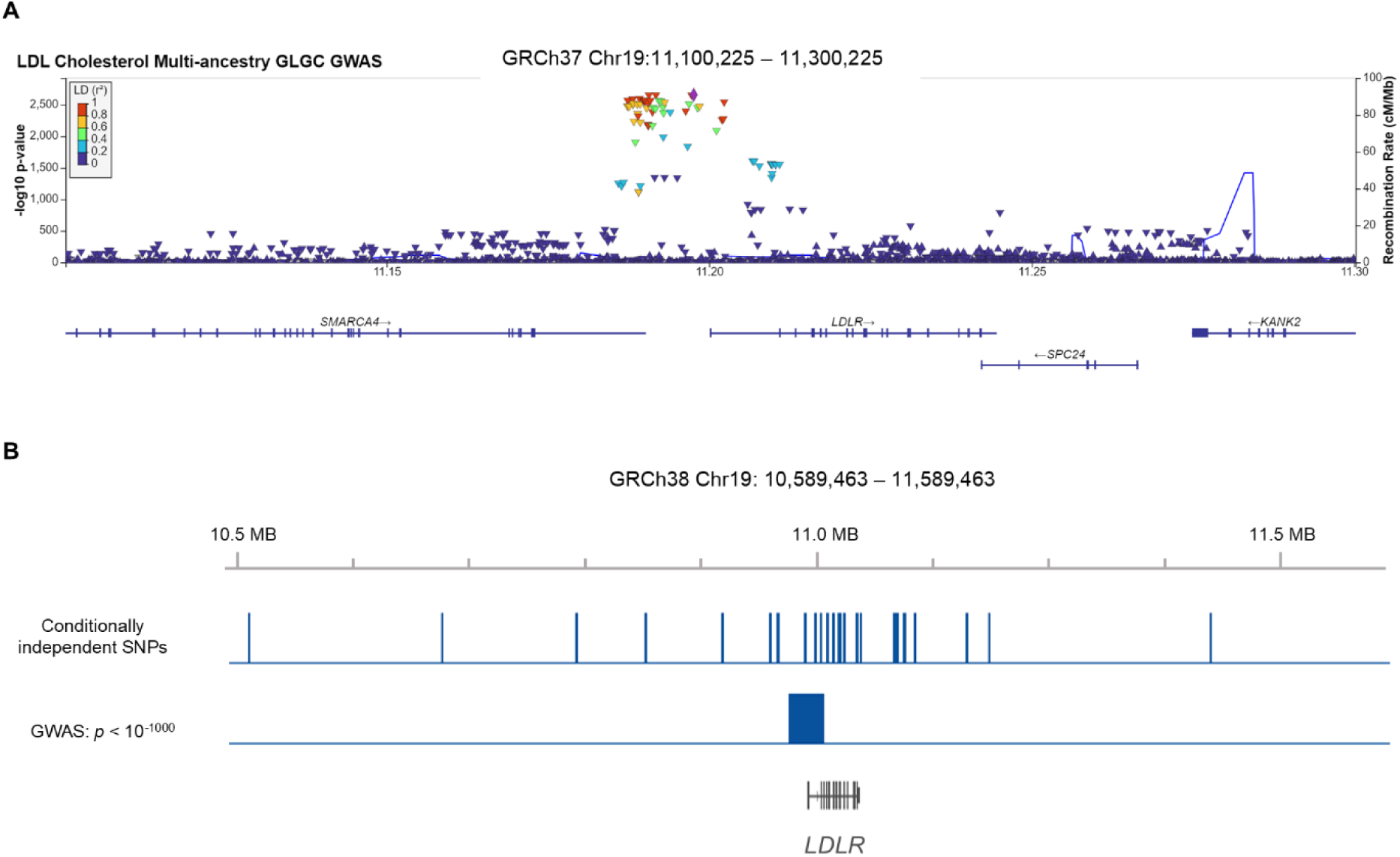
LDL cholesterol-associated variants at the *LDLR* locus. (A) LocusZoom plot of *LDLR* variant associations with LDL cholesterol, with individual SNPs shaded by their linkage disequilibrium with the lead SNP. (B) Genomic tracks of regions targeted by the CRISPR library, including a block containing all variants identified in (A) and 1 kb intervals centered on other SNPs within 500 kb of the *LDLR* TSS and an independent association with LDL cholesterol after conditioning on the lead SNP. Source data is provided in Supplemental Tables 1 and 2.

**Figure S2.**
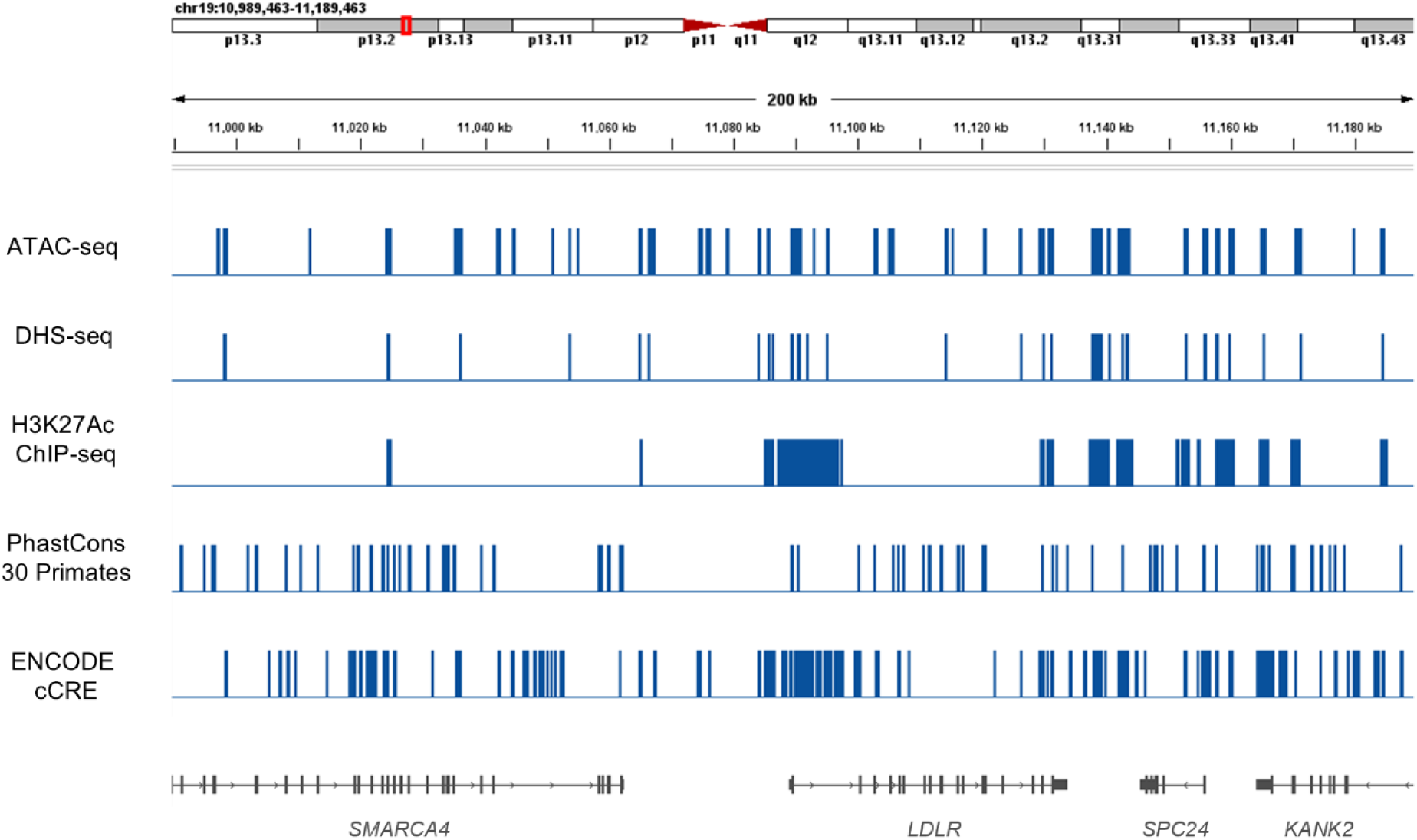
Candidate CRE nomination. Genomic tracks of CRISPR library target regions selected based on the indicated supporting information, as described in Materials and Methods.

**Figure S3.**
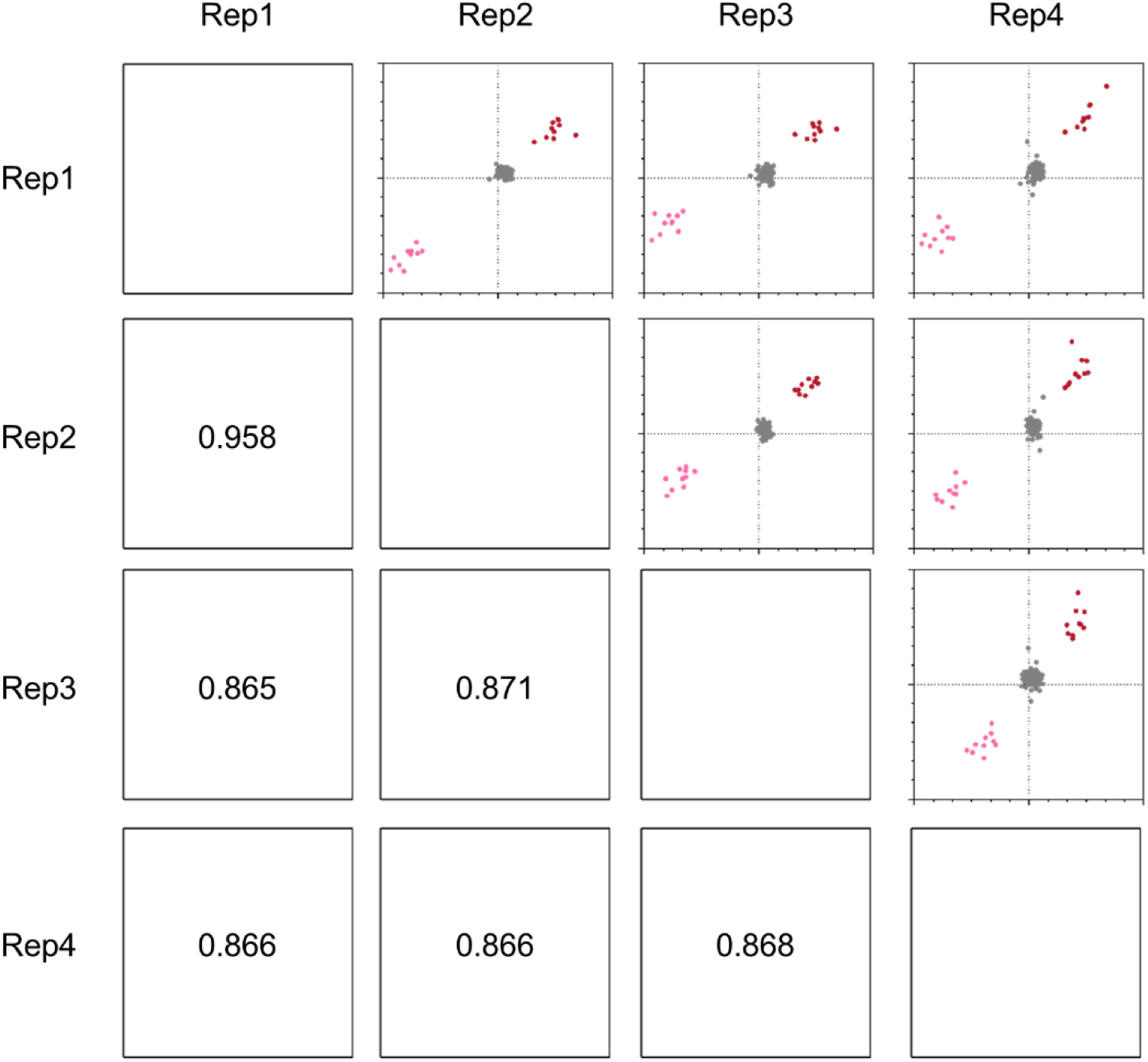
Reproducibility of control gRNA segregation. Pairwise correlations between the log2 fold change of individual control gRNA abundance in LDL^high^ vs LDL^low^ cells for independent replicates of the CRISPR screen. Dark red data points correspond to gRNAs expected to be enriched in LDL^high^ cells due to their disruption of the *MYLIP* coding sequence. Pink data points correspond to gRNAs expected to be depleted in LDL^high^ cells due to their disruption of the *LDLR* coding sequence. Gray data points correspond to gRNAs expected to have no effect on LDL uptake related to their targeting of the AAVS safe harbor region.

**Figure S4.**
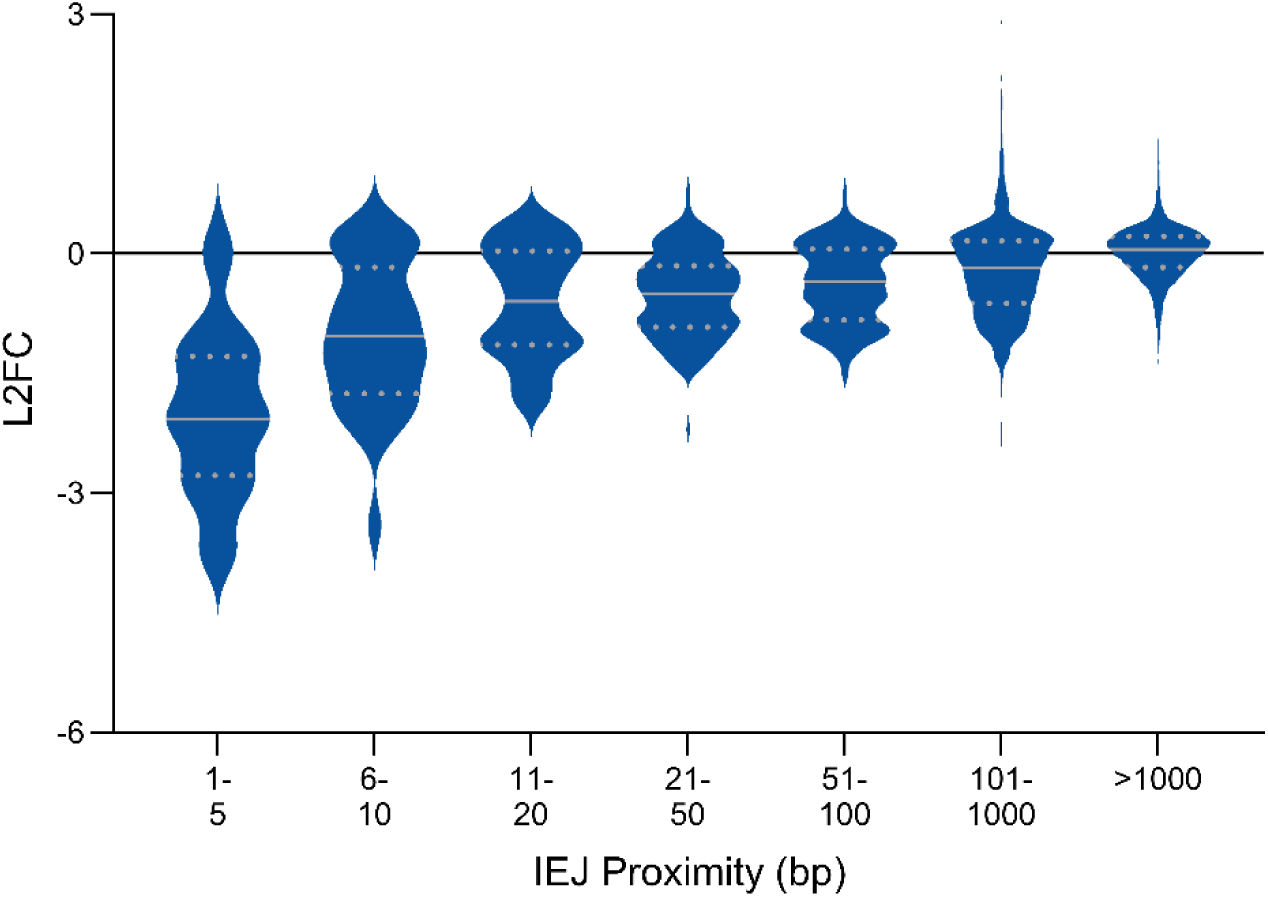
Relationship of gRNA enrichment to proximity to intron-exon junctions. Violin plots for average log2 fold-change gRNA abundance in LDL^high^ relative to LDL^low^ cells for individual gRNAs grouped by their proximity to the nearest *LDLR* intron-exon junction. Solid and dashed lines represent the median and quartiles, respectively, for each group.

**Figure S5.**
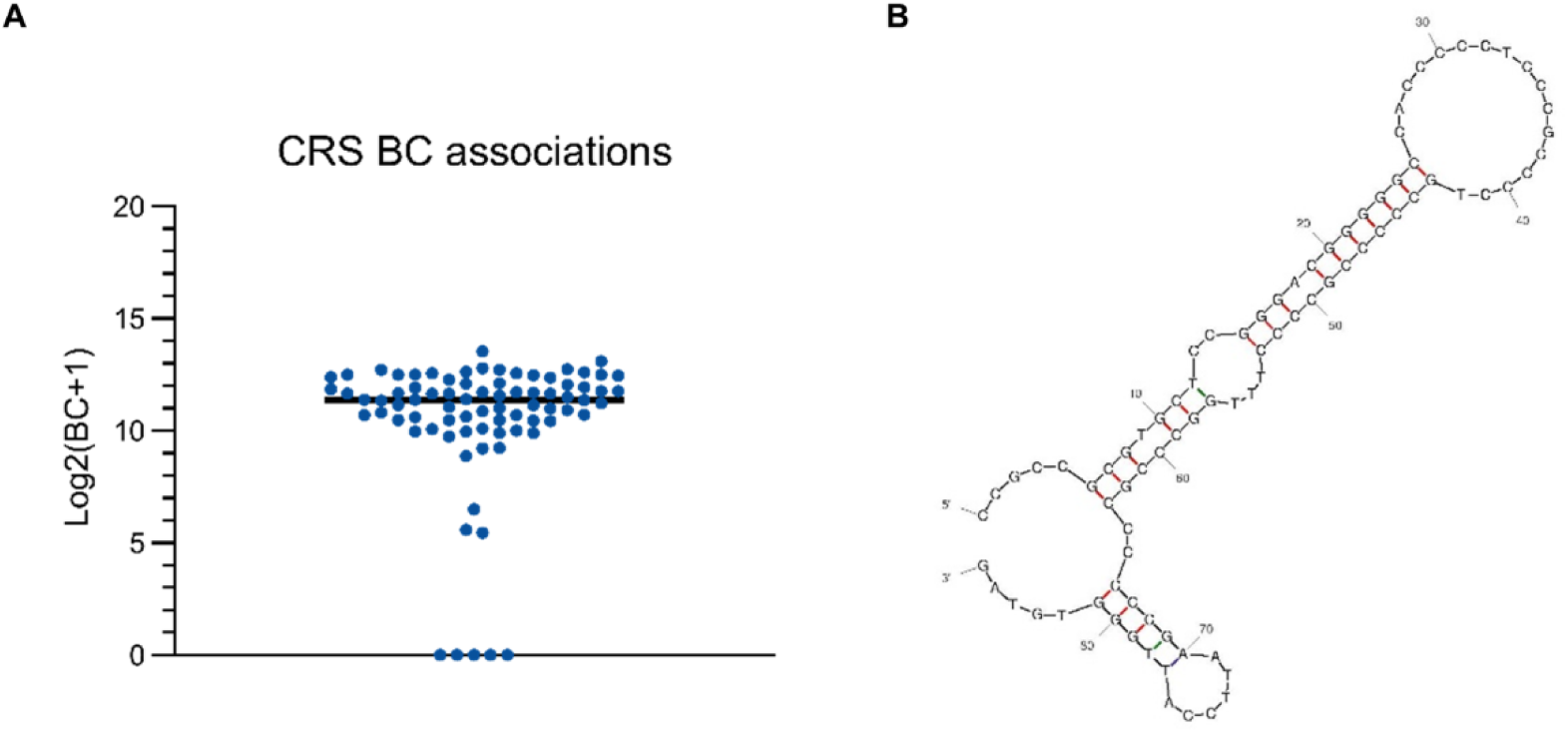
Analysis of MPRA library synthesis. (A) The number of individual 5’ UTR barcodes detected in association with each expected *cis*-regulatory sequence in the MPRA library. Source data is provided in Supplemental Table 6. (B) Predicted hairpin structure in the sequence common to all expected *cis*-regulatory sequences absent from the plasmid library.

**Figure S6.**
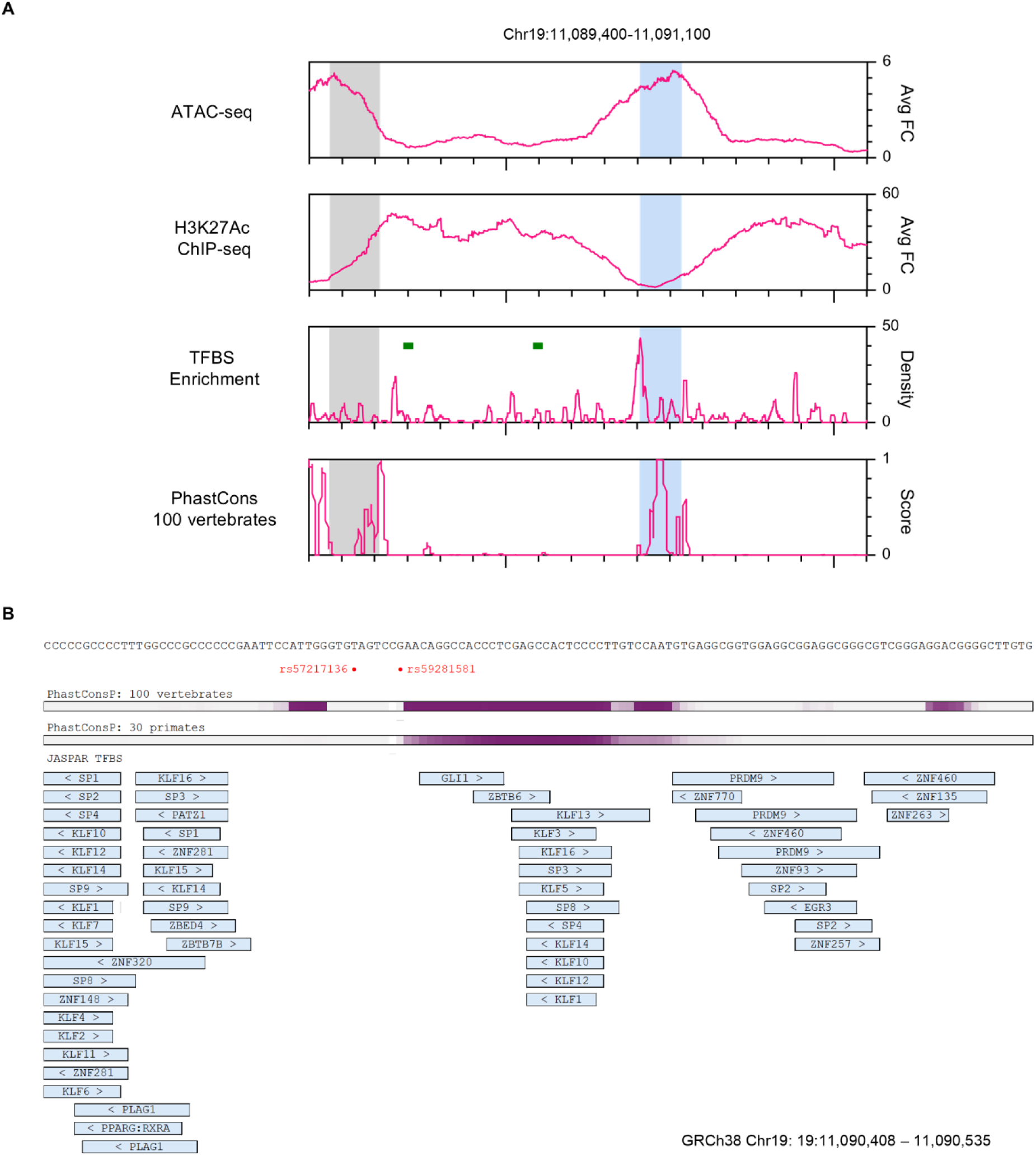
Features of the fine-mapped first intronic *LDLR* enhancer. (A) Genomic tracks from Fig 2 overlaid with light blue shading to indicate the position of the MPRA fine-mapped enhancer interval. (B) Genomic sequence of the fine-mapped interval aligned with positions of human genetic variants of interest, a heat map of PhasConsP evolutionary conservation scores among 100 vertebrates and 30 primates (deeper shades of purple indicate greater conservation), and transcription factor binding motifs identified by JASPAR.

**Supplemental Table 1. LDL cholesterol GWAS associations.** Common genetic variants at the *LDLR* locus associated with LDL cholesterol at a significance threshold of *p* < 10^-1000^ in Graham et al^14^.

**Supplemental Table 2. Conditional analysis of LDL cholesterol-associated *LDLR* variants.** LDL cholesterol associations derived from Graham et al^14^ for individuals of European ancestry before and after adjustment for linkage disequilibrium with the lead SNP (rs73015024). All noncoding variants within 500 kb of the *LDLR* TSS with an allele frequency greater than 1% were analyzed; those with significant associations after conditional analysis are displayed.

**Supplemental Table 3. CRISPR library composition.** Identifier and sequences for every gRNA in the CRISPR library used in this study. For gRNAs targeting potential *LDLR cis*-regulatory sequences, the genomic coordinate for its predicted double-strand break and the basis for its selection are also provided.

**Supplemental Table 4. CRISPR screen results.** DESeq2 output for the relative change in abundance of each indicated gRNA in LDL^high^ versus LDL^low^ cells.

**Supplemental Table 5. MPRA library design.** Genomic coordinates and sequences for each indicated tile in the MPRA library. The test sequence constitutes the major allele at each position as determined by population allele frequences in Gnomad, notably including the presence of the common insertion rs59281581 that is not present in the hg38 human genome reference sequence.

**Supplemental Table 6. Association of 5’ UTR barcodes with *cis*-regulatory sequences.** The number of reads for each combination of 5’ UTR barcode sequence and CRS test sequence are listed together with the overall share of reads for a given 5’ UTR barcode that match the indicated CRS. Filtering was performed to remove any combination of 5’ UTR barcode and CRS with less than 10 reads, or any 5’ UTR barcode with less than 99% of reads associated with the same CRS.

**Supplemental Table 7. MPRA expression data.** Aggregate counts per million for the 5’ UTR barcodes corresponding to each *cis*-regulatory sequence in the MPRA library, demultiplexed by sample identity.

**Supplemental Table 8. MPRA expression analysis.** Expression associated with each CRS, as quantified by the ratio of aggregate counts for its associated 5’UTR barcodes in cDNA relative to genomic DNA, for each of 5 independent replicates in HuH7 and HepG2 cells. Ratios are normalized to the average values for negative control sequences for each sample.

**Supplemental Table 9. Oligonucleotides.** Sequences for the primers used in this study are provided and grouped by application.

**Supplemental Table 10. Source data for *LDLR* candidate CRE nomination.** Identifying information for the ENCODE data sets used in the design of the custom CRISPR library for this study.

## REFERENCES

1 Cohen, J. C., Boerwinkle, E., Mosley, T. H., Jr. & Hobbs, H. H. Sequence variations in PCSK9, low LDL, and protection against coronary heart disease. N Engl J Med 354, 1264–1272, doi:10.1056/NEJMoa054013 (2006).

2 Musunuru, K. & Kathiresan, S. Genetics of Common, Complex Coronary Artery Disease. Cell 177, 132–145, doi:10.1016/j.cell.2019.02.015 (2019).

3 Boren, J. et al. Low-density lipoproteins cause atherosclerotic cardiovascular disease: pathophysiological, genetic, and therapeutic insights: a consensus statement from the European Atherosclerosis Society Consensus Panel. Eur Heart J 41, 2313–2330, doi:10.1093/eurheartj/ehz962 (2020).

4 Horton, J. D., Goldstein, J. L. & Brown, M. S. SREBPs: activators of the complete program of cholesterol and fatty acid synthesis in the liver. J Clin Invest 109, 1125–1131, doi:10.1172/JCI15593 (2002).

5 Briggs, M. R., Yokoyama, C., Wang, X., Brown, M. S. & Goldstein, J. L. Nuclear protein that binds sterol regulatory element of low density lipoprotein receptor promoter. I. Identification of the protein and delineation of its target nucleotide sequence. J Biol Chem 268, 14490–14496 (1993).

6 Shimano, H. Sterol regulatory element-binding proteins (SREBPs): transcriptional regulators of lipid synthetic genes. Prog Lipid Res 40, 439–452, doi:10.1016/s0163-7827(01)00010-8 (2001).

7 Gasperini, M., Tome, J. M. & Shendure, J. Towards a comprehensive catalogue of validated and target-linked human enhancers. Nat Rev Genet 21, 292–310, doi:10.1038/s41576-019-0209-0 (2020).

8 Kong, W. J., Liu, J. & Jiang, J. D. Human low-density lipoprotein receptor gene and its regulation. J Mol Med (Berl) 84, 29–36, doi:10.1007/s00109-005-0717-6 (2006).

9 Emmer, B. T. et al. Genome-scale CRISPR screening for modifiers of cellular LDL uptake. PLoS Genet 17, e1009285, doi:10.1371/journal.pgen.1009285 (2021).

10 Smith, G. A. et al. Cold shock domain-containing protein E1 is a posttranscriptional regulator of the LDL receptor. Sci Transl Med 14, eabj8670, doi:10.1126/scitranslmed.abj8670 (2022).

11 Hamilton, M. C. et al. Systematic elucidation of genetic mechanisms underlying cholesterol uptake. Cell Genom 3, 100304, doi:10.1016/j.xgen.2023.100304 (2023).

12 Willer, C. J. et al. Discovery and refinement of loci associated with lipid levels. Nat Genet 45, 1274–1283, doi:10.1038/ng.2797 (2013).

13 Buniello, A. et al. The NHGRI-EBI GWAS Catalog of published genome-wide association studies, targeted arrays and summary statistics 2019. Nucleic Acids Res 47, D1005–D1012, doi:10.1093/nar/gky1120 (2019).

14 Graham, S. E. et al. The power of genetic diversity in genome-wide association studies of lipids. Nature 600, 675–679, doi:10.1038/s41586-021-04064-3 (2021).

15 Fairoozy, R. H., White, J., Palmen, J., Kalea, A. Z. & Humphries, S. E. Identification of the Functional Variant(s) that Explain the Low-Density Lipoprotein Receptor (LDLR) GWAS SNP rs6511720 Association with Lower LDL-C and Risk of CHD. PLoS One 11, e0167676, doi:10.1371/journal.pone.0167676 (2016).

16 Consortium, E. P. An integrated encyclopedia of DNA elements in the human genome. Nature 489, 57–74, doi:10.1038/nature11247 (2012).

17 Meuleman, W. et al. Index and biological spectrum of human DNase I hypersensitive sites. Nature 584, 244–251, doi:10.1038/s41586-020-2559-3 (2020).

18 Vierstra, J. et al. Global reference mapping of human transcription factor footprints. Nature 583, 729–736, doi:10.1038/s41586-020-2528-x (2020).

19 Sethi, A. et al. Supervised enhancer prediction with epigenetic pattern recognition and targeted validation. Nat Methods 17, 807–814, doi:10.1038/s41592-020-0907-8 (2020).

20 Khan, T. G., Ginsburg, D. & Emmer, B. T. The small GTPase RAB10 regulates endosomal recycling of the LDL receptor and transferrin receptor in hepatocytes. J Lipid Res 63, 100248, doi:10.1016/j.jlr.2022.100248 (2022).

21 Serrano-Zayas, C. et al. Regulation of cellular LDL uptake by PROX1 and CHD7. bioRxiv, 2022.2009.2020.507601, doi:10.1101/2022.09.20.507601 (2022).

22 Consortium, E. P. et al. Expanded encyclopaedias of DNA elements in the human and mouse genomes. Nature 583, 699–710, doi:10.1038/s41586-020-2493-4 (2020).

23 Papapetrou, E. P. & Schambach, A. Gene Insertion Into Genomic Safe Harbors for Human Gene Therapy. Mol Ther 24, 678–684, doi:10.1038/mt.2016.38 (2016).

24 Blattmann, P. et al. Systems Pharmacology Dissection of Cholesterol Regulation Reveals Determinants of Large Pharmacodynamic Variability between Cell Lines. Cell Syst 5, 604–619 e607, doi:10.1016/j.cels.2017.11.002 (2017).

25 Pundhir, S., Bagger, F. O., Lauridsen, F. B., Rapin, N. & Porse, B. T. Peak-valley-peak pattern of histone modifications delineates active regulatory elements and their directionality. Nucleic Acids Res 44, 4037–4051, doi:10.1093/nar/gkw250 (2016).

26 Castro-Mondragon, J. A. et al. JASPAR 2022: the 9th release of the open-access database of transcription factor binding profiles. Nucleic Acids Res 50, D165–D173, doi:10.1093/nar/gkab1113 (2022).

27 Gordon, M. G. et al. lentiMPRA and MPRAflow for high-throughput functional characterization of gene regulatory elements. Nat Protoc 15, 2387–2412, doi:10.1038/s41596-020-0333-5 (2020).

28 Banerji, J., Rusconi, S. & Schaffner, W. Expression of a beta-globin gene is enhanced by remote SV40 DNA sequences. Cell 27, 299–308, doi:10.1016/0092-8674(81)90413-x (1981).

29 Notani, D. First glimpse of enhancers in gene regulation. Nat Rev Genet 23, 522–523, doi:10.1038/s41576-022-00492-7 (2022).

30 Inoue, F. & Ahituv, N. Decoding enhancers using massively parallel reporter assays. Genomics 106, 159–164, doi:10.1016/j.ygeno.2015.06.005 (2015).

31 Chen, S. et al. A genomic mutational constraint map using variation in 76,156 human genomes. Nature 625, 92–100, doi:10.1038/s41586-023-06045-0 (2024).

32 Klein, J. C. et al. A systematic evaluation of the design and context dependencies of massively parallel reporter assays. Nat Methods 17, 1083–1091, doi:10.1038/s41592-020-0965-y (2020).

33 Moreau, P. et al. The SV40 72 base repair repeat has a striking effect on gene expression both in SV40 and other chimeric recombinants. Nucleic Acids Res 9, 6047–6068, doi:10.1093/nar/9.22.6047 (1981).

34 Wang, X. et al. High-resolution genome-wide functional dissection of transcriptional regulatory regions and nucleotides in human. Nat Commun 9, 5380, doi:10.1038/s41467-018-07746-1 (2018).

35 Shlyueva, D., Stampfel, G. & Stark, A. Transcriptional enhancers: from properties to genome-wide predictions. Nat Rev Genet 15, 272–286, doi:10.1038/nrg3682 (2014).

36 Zaugg, J. B. et al. Current challenges in understanding the role of enhancers in disease. Nat Struct Mol Biol 29, 1148–1158, doi:10.1038/s41594-022-00896-3 (2022).

37 Kwasnieski, J. C., Fiore, C., Chaudhari, H. G. & Cohen, B. A. High-throughput functional testing of ENCODE segmentation predictions. Genome Res 24, 1595–1602, doi:10.1101/gr.173518.114 (2014).

38 Montalbano, A., Canver, M. C. & Sanjana, N. E. High-Throughput Approaches to Pinpoint Function within the Noncoding Genome. Mol Cell 68, 44–59, doi:10.1016/j.molcel.2017.09.017 (2017).

39 Canver, M. C. et al. BCL11A enhancer dissection by Cas9-mediated in situ saturating mutagenesis. Nature 527, 192–197, doi:10.1038/nature15521 (2015).

40 Sanjana, N. E. et al. High-resolution interrogation of functional elements in the noncoding genome. Science 353, 1545–1549, doi:10.1126/science.aaf7613 (2016).

41 Diao, Y. et al. A new class of temporarily phenotypic enhancers identified by CRISPR/Cas9-mediated genetic screening. Genome Res 26, 397–405, doi:10.1101/gr.197152.115 (2016).

42 Rajagopal, N. et al. High-throughput mapping of regulatory DNA. Nat Biotechnol 34, 167–174, doi:10.1038/nbt.3468 (2016).

43 Sen, D. R. et al. The epigenetic landscape of T cell exhaustion. Science 354, 1165–1169, doi:10.1126/science.aae0491 (2016).

44 Pollak, N. M., Hoffman, M., Goldberg, I. J. & Drosatos, K. Kruppel-like factors: Crippling and un-crippling metabolic pathways. JACC Basic Transl Sci 3, 132–156, doi:10.1016/j.jacbts.2017.09.001 (2018).

45 Alaiti, M. A., Orasanu, G., Tugal, D., Lu, Y. & Jain, M. K. Kruppel-like factors and vascular inflammation: implications for atherosclerosis. Curr Atheroscler Rep 14, 438–449, doi:10.1007/s11883-012-0268-6 (2012).

46 Sanchez, H. B., Yieh, L. & Osborne, T. F. Cooperation by sterol regulatory element-binding protein and Sp1 in sterol regulation of low density lipoprotein receptor gene. J Biol Chem 270, 1161–1169, doi:10.1074/jbc.270.3.1161 (1995).

47 Dawson, P. A. et al. Sterol-dependent repression of low density lipoprotein receptor promoter mediated by 16-base pair sequence adjacent to binding site for transcription factor Sp1. J Biol Chem 263, 3372–3379 (1988).

48 Hui, C. C. & Angers, S. Gli proteins in development and disease. Annu Rev Cell Dev Biol 27, 513–537, doi:10.1146/annurev-cellbio-092910-154048 (2011).

49 Sanjana, N. E., Shalem, O. & Zhang, F. Improved vectors and genome-wide libraries for CRISPR screening. Nat Methods 11, 783–784, doi:10.1038/nmeth.3047 (2014).

50 Ran, F. A. et al. Genome engineering using the CRISPR-Cas9 system. Nat Protoc 8, 2281–2308, doi:10.1038/nprot.2013.143 (2013).

51 Emmer, B. T. et al. The cargo receptor SURF4 promotes the efficient cellular secretion of PCSK9. Elife 7, doi:10.7554/eLife.38839 (2018).

52 Graham, S. E. et al. The power of genetic diversity in genome-wide association studies of lipids. Nature 600, 675–679, doi:10.1038/s41586-021-04064-3 (2021).

53 Boughton, A. P. et al. LocusZoom.js: interactive and embeddable visualization of genetic association study results. Bioinformatics 37, 3017–3018, doi:10.1093/bioinformatics/btab186 (2021).

54 Yang, J. et al. Conditional and joint multiple-SNP analysis of GWAS summary statistics identifies additional variants influencing complex traits. Nat Genet 44, 369–375, S361–363, doi:10.1038/ng.2213 (2012).

55 Karolchik, D. et al. The UCSC Table Browser data retrieval tool. Nucleic Acids Res 32, D493–496, doi:10.1093/nar/gkh103 (2004).

56 Murphy, W. J. et al. Resolution of the early placental mammal radiation using Bayesian phylogenetics. Science 294, 2348–2351, doi:10.1126/science.1067179 (2001).

57 Sherman, E. J. et al. Identification of cell type specific ACE2 modifiers by CRISPR screening. PLoS Pathog 18, e1010377, doi:10.1371/journal.ppat.1010377 (2022).

58 Bragazzi Cunha, J., et al. Type I interferon signaling induces a delayed antiproliferative response in respiratory epithelial cells during SARS-CoV-2 infection. J Virol 97, e0127623, doi:10.1128/jvi.01276-23 (2023).

59 Li, W. et al. MAGeCK enables robust identification of essential genes from genome-scale CRISPR/Cas9 knockout screens. Genome Biol 15, 554, doi:10.1186/s13059-014-0554-4 (2014).

60 Zuker, M. Mfold web server for nucleic acid folding and hybridization prediction. Nucleic Acids Res 31, 3406–3415, doi:10.1093/nar/gkg595 (2003).

61 Robinson, J. T. et al. Integrative genomics viewer. Nat Biotechnol 29, 24–26, doi:10.1038/nbt.1754 (2011).

